# Albatrosses employ orientation and routing strategies similar to yacht racers

**DOI:** 10.1101/2022.12.13.520348

**Authors:** Yusuke Goto, Henri Weimerskirch, Keiichi Fukaya, Ken Yoda, Masaru Naruoka, Katsufumi Sato

## Abstract

The way goal-oriented birds adjust their travel direction and route in response to wind significantly affects their travel costs. This is expected to be particularly pronounced in pelagic seabirds, which utilize a wind-dependent flight style called dynamic soaring. Dynamic soaring seabirds in situations without a definite goal, e.g. searching for prey, are known to preferentially fly with tail-to-side winds to increase the speed and search area, and reduce travel costs. However, little is known about their reaction to wind when heading to a definite goal, such as homing. Homing tracks of wandering albatrosses (*Diomedea exulans*) vary from beelines to zigzags, which are similar to those of sailboats. Here, given that both albatrosses and sailboats travel slower in headwinds and tailwinds, we tested whether the time-minimizing strategies used by yacht racers can be compared to the locomotion patterns of wandering albatrosses. We predicted that when the goal is located upwind or downwind, albatrosses should deviate their travel directions from the goal on the microscale and increase the number of turns on the macroscale. Both hypotheses were supported by track data from albatrosses and racing yachts in the Southern Ocean confirming that albatrosses qualitatively employ the same strategy as yacht racers. Nevertheless, albatrosses did not strictly minimize their travel time, likely making their flight robust against wind fluctuations to reduce flight costs. Our study provides the first empirical evidence of tacking in albatrosses and demonstrates that man-made movement strategies provide a new perspective on the laws underlying wildlife movement.

## Introduction

Birds traverse great distances to reach their goals. Some species migrate across the globe from their wintering grounds to breeding grounds (1, 2), and some species undertake long-distance foraging trips during the breeding season, traveling hundreds or thousands of kilometers from their nests, and then returning to feed their chicks (3–9). These long-distance flights require significant travel costs, such as energy and time. Wind significantly impacts these cost requirements: tailwinds increase birds’ speed, headwinds slow them down, and crosswinds can divert them off course (6, 10, 11). Therefore, through natural selection, birds are expected to have acquired a navigational capacity that allows them to select routes that reduce travel costs under wind conditions they encounter (12–14). This macro-scale route selection consists of a series of decisions related to travel direction, taking into account the wind and goal directions, a process referred to as ’orientation’ (11). Orientation is likely determined by micro-scale flight dynamics, specifically, the variations in energy and time needed to travel a given distance, which depends on the travel direction relative to the wind and goal directions (15, 16). Accordingly, to gain a deeper understanding of birds’ hierarchically structured navigation under wind conditions, we need to predict their orientation and route selection based on micro-scale flight dynamics and compare these predictions with real data (16). However, while bird orientation and route selection in wind conditions have been extensively studied at a macro scale, empirical research on how micro-scale flight dynamics shape these aspects is still limited.

Among various bird species, procellariiform seabirds (i.e., petrels, shearwaters, and albatrosses) are particularly distinctive due to their unique wind-utilizing flight style, and may provide a valuable opportunity to study the implications of micro-scale flight dynamics on orientation and route selection. During their breeding period, these seabirds can fly hundreds of kilometers away from their nests (3–9). During their foraging trips, birds spend only a small fraction of their flight time flapping their wings. For example, wandering albatrosses (*Diomedea exulans*), one of the largest dynamic soaring species, only spend 1–15% of their flight time flapping their wings (17). This efficient travel is facilitated by a flight style known as dynamic soaring, where the birds exploit mechanical energy from the atmosphere by utilizing wind speeds that increase with the altitude above the sea surface (18–20). Although this wind-dependent flight mechanism, which influences travel speed and energy consumption rate (21–23), could affect orientation and route selection, these aspects remain largely unexplored in spite of numerous tracking studies on procellariiform seabirds conducted over the last three decades (3, 24, 25). Previous studies have demonstrated that procellariiform seabirds tend to prefer tail-to-side winds during their foraging trips (8, 9, 21, 26–30). However, it is important to note that these studies mainly focused on travel and wind direction, neglecting a crucial factor for understanding birds’ navigation strategies—the direction of their goal (11).

The significance of the goal direction during the foraging trips of pelagic seabirds changes with each phase of the trip. As same as other central place foragers (31), foraging trips of pelagic seabirds are often categorized into three phases: outbound, middle, and returning (also referred to as the homing phase) (29). Importantly, the factors affecting the locomotion decision-making of pelagic seabirds vary between the non-homing phases (outbound and middle) and the homing phase. During the non-homing phases, their goal locations, if any, typically encompass large areas like a frontal zone (32), spanning several hundred kilometers. Consequently, the necessity to reach a specific destination is weak or non-existent, leading birds to select a travel direction that maximizes distance per unit of travel cost. Therefore dynamic soaring seabirds are expected to prefer tail-to-side winds that enable higher travel speed and lower energy costs (21) as consistent with previous studies (8, 9, 21, 26–30). In contrast, during the homing phase, the nest serves as a distinct goal. Therefore, a conflict may arise between the alignment to the goal direction and the birds’ preference for sidewinds, especially when the goal is not located in the crosswind direction. Hence, it’s essential to study their homing tracks under various wind and goal directions to understand how micro-scale locomotion dynamics (in this case, dynamic soaring) affect animal orientation and route selection—a topic that has not been extensively explored. The diverse homing tracks of wandering albatrosses may serve as a distinctive illustration of this unresolved issue. Fig. 1 illustrates the homing portion of the foraging trip of the wandering albatross, which we arbitrarily selected from previously published track data (33, 34). Their homing tracks exhibit a variety of patterns, ranging from straight lines to zigzags, similar to the tracks of sailboats (Fig. 1). This leads to a question: is the similarity between albatross and sailboat trajectories merely artificial, resulting from our selective choice of tracks, or does a common underlying rule govern their patterns, possibly their response to wind?

**Fig. 1.**
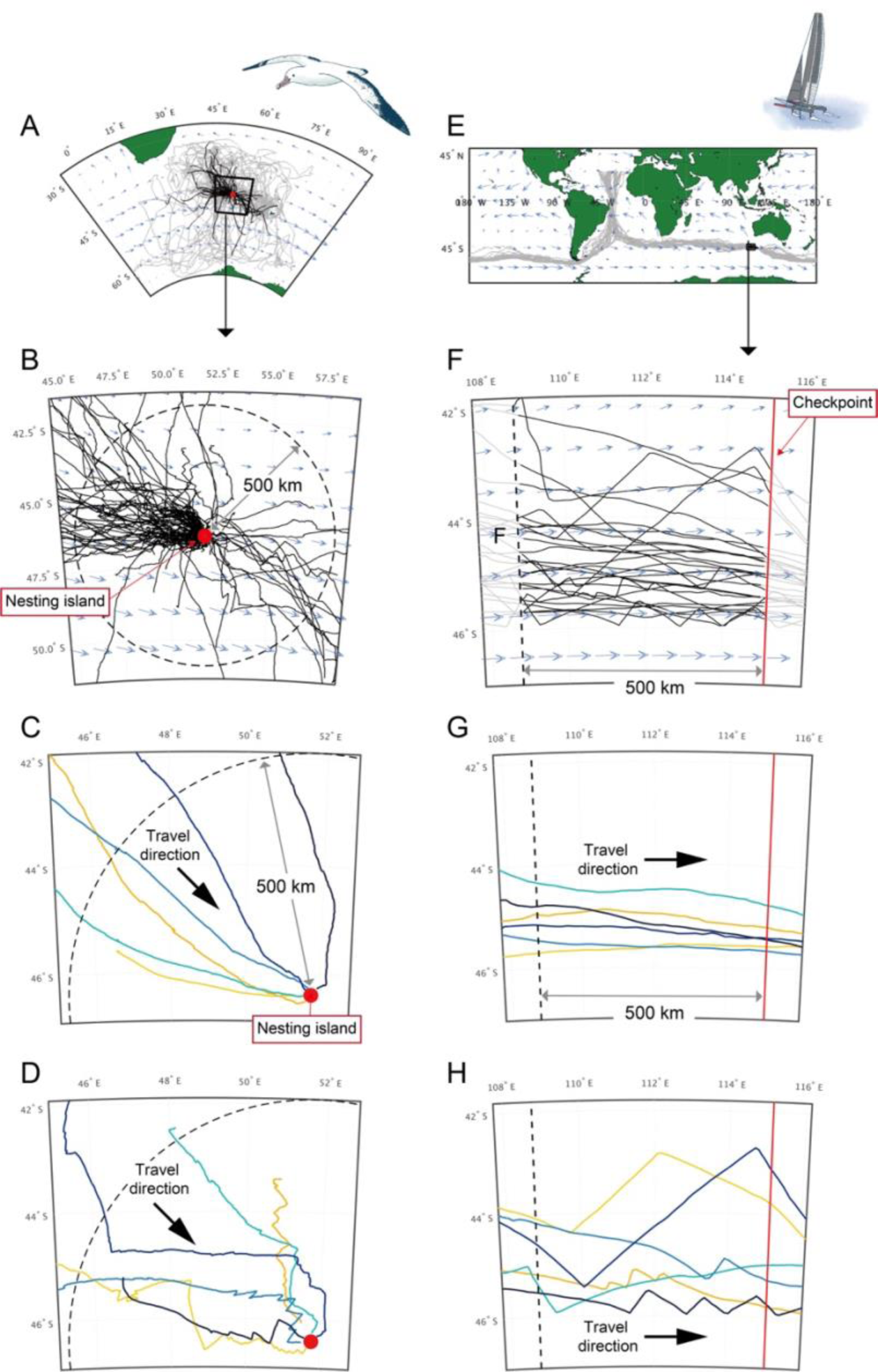
Tracks of albatrosses and racing sailboats. (A) Foraging trips of wandering albatrosses recorded by GPS (*N* = 149). Portions of tracks in homing phase are shown in black lines and the other portions are shown in grey lines. (B) Homing tracks of albatrosses within 500 km from the nesting island (Possession Island). Light blue arrows in (A–B) represent average winds for January 2018 based on ERA5 ECMWF. (C–D) Homing tracks of albatrosses in straight lines (C) and winding patterns (D) selected from trajectories shown in (B). Differences in color represent individuals. (E) Tracks of sailboats in the 2020 Vendée Globe race around the world. (F) Tracks (black lines) of sailboats within 500 km from the middle checkpoint (red line). Tracks in straight lines (G) and winding patterns (H), selected from trajectories shown in (F). Light blue arrows in (E–F) represent average winds for December 2020 based on ERA5 ECMWF. (G–H). Differences in color represent each sailboat.

Recent studies (20, 23) have highlighted the similarities between the mechanisms of dynamic soaring and sailing, especially regarding how wind direction impacts the travel speeds of sailboats and birds, which are maximized in crosswinds and reduced in tailwinds and headwinds. As the speed of a sailboat depends on wind direction, yacht racers often adjust their course away from the goal if it lies leeward or windward. They alternatively switch their travel direction, known as ’tacking’, which results in a zigzag route on a macro scale (35, 36). This approach enables them to reach the goal more quickly. Given the wind-dependent speed similarity between dynamic soaring birds and sailboats, here we hypothesize that albatrosses adopt navigation strategies similar to yacht racers during homing to minimize travel time and energy consumption reaching their nests.

Here, we tested whether the time-minimizing strategies used by yacht racers can explain the orientation and route selection of wandering albatrosses. We predicted that when the goal is located upwind or downwind, albatrosses should (i) deviate their travel directions from the goal at the microscale and (ii) increase the number of turns at the macroscale (see *prediction of albatross movement based on time-minimizing orientation strategy by sailors* in **Results**). We tested these predictions with tracking data from albatrosses and racing yachts in the Southern Ocean. Testing these predictions poses a challenge as it involves a detailed examination of the travel directions of homing albatrosses under various wind and goal conditions, necessitating substantial high-resolution tracking data. To address this, we utilized a comprehensive set of tracking data from their foraging trips, which comprises 149 tracks or a total of 407,659 fix data points, each recorded at a fix point every two minutes.

## Results

First, we confirmed the similarities in wind-dependent travel speeds between sailboats and albatrosses (23). Thereafter, we derived predictions of albatross movement based on the time-minimizing orientation strategies used by yacht racers. Then, we tested these predictions qualitatively and quantitatively by using track data of albatross and sailboats.

### Similarities in wind-dependent travel speeds between sailboats and albatrosses

We analyzed track data (1 data point every 2 min) from 149 foraging trips made by wandering albatrosses during their breeding period from Possession Island, Crozet Islands, and track data (1 data point every 30 min) from 28 yachts participating in the 2020 ‘Vendée Globe’, a non-stop round-the-world yacht race across the Southern Ocean (Fig. 1). Both albatrosses and sailboats mainly traveled in 40–60°S latitudes and were constantly exposed to strong winds (average wind speed was 8.7 ± 3.4 m s^−1^ for albatross and 8.1 ± 3.0 m s^−1^ for sailboats).

We calculated the speed and direction of the albatross and sailboats by computing a vector connecting two successive data points (*N* = 407,659 for albatrosses and *N* = 102,922 for sailboats); the average distances traveled between the two observation points by the albatrosses and sailboats were 1.5 ± 0.6 km and 12.1 ± 3.4 km, respectively. Consequently, the average travel speed was 12.4 ± 5.2 m s^−1^ for albatrosses and 6.7 ± 1.9 m s^−1^ for sailboats. Their travel speed changed according to the travel direction relative to the wind direction (Fig. 2). Such plots are called ‘polar diagrams’ in the field of sailing. In theory, the speed of sailboats decreases in tailwinds and headwinds, thereby creating butterfly-shaped polar diagrams (35). A recent study reported that the polar diagrams of wandering albatrosses were also butterfly-shaped, based on track data at around 1 h sampling intervals (23). We confirmed the butterfly-shaped polar diagrams of the sailboats and albatrosses in our data (2 min sampling intervals) by fitting non-parametric functions (see **METHODS**). As described below, these butterfly-shaped polar diagrams influence the movement strategies of yacht racers and are also expected to influence those of albatrosses.

**Fig. 2.**
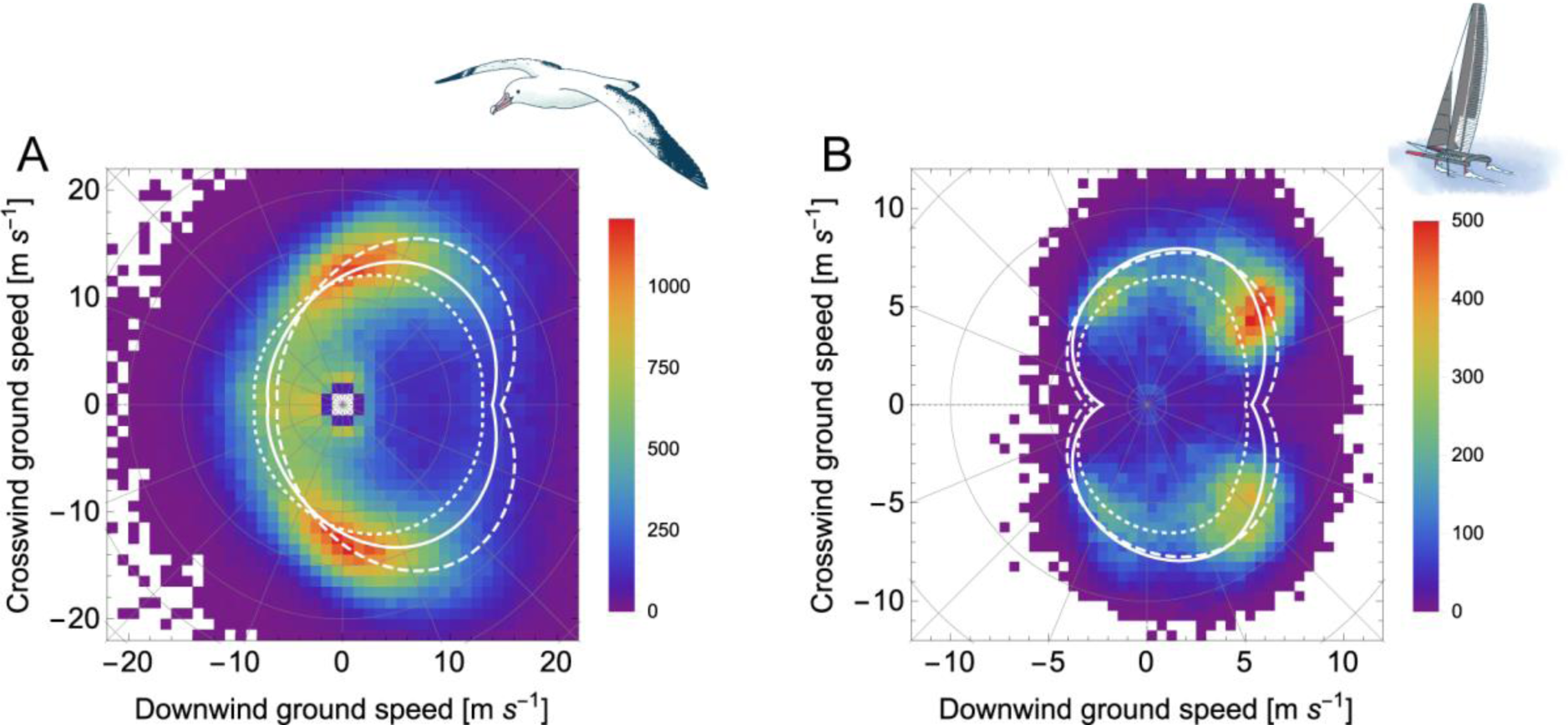
Similarities in wind-dependent travel speeds between sailboats and albatrosses. Two-dimensional histogram of the ground velocity of albatrosses (A) and sailboats (B). Histograms in bins of 1 m s^−1^ for albatrosses and 0.5 m s^−1^ for sailboats. Colors represent the number of data points in each bin. The white lines indicate polar diagrams obtained by applying the general additive models (GAM) to the data. The solid line shows the polar diagrams with a 9 m s^−1^ wind speed, dotted lines with 6 m s^−1^ wind speed, and dashed lines with 12 m s^−1^ wind speed.

### Prediction of albatross movement based on time-minimizing orientation strategy by sailors

We examine situations where both sailboat racers and albatrosses have goals: racers aim for an intermediate checkpoint, while albatrosses aim to return to their nesting island (Fig. 1). Several factors can influence their decision-making: the spatiotemporal pattern of wind conditions en route to the goal, wind-influenced speed (23), wind-dependent energy expenditure (21), and the spatial range within which they can perceive or predict wind conditions. In this study, we make two assumptions: 1) both albatrosses and yacht racers choose their travel direction based on the local wind conditions where they are, and 2) their objective is to minimize travel time to reach their goals. The second assumption is evident in yacht racing, but we argue it is also applicable to albatrosses, for two reasons. Firstly, by minimizing travel time, albatrosses can return to their nests sooner, thereby reducing the risk of their partners abandoning the nests, which protects the eggs or chicks (37, 38). Secondly, decreasing travel time leads to reduced energy expenditure, which benefits the survival and reproductive success of the albatrosses (39). If the dependency of energy consumption rate on the travel direction relative to the wind can be ignored, then the energy required to cover a given distance is proportionate to the elapsed travel time. Hence, elapsed time can serve as an effective proxy for energy consumption (this point will be revisited in **Discussion**).

Based on these assumptions, we derived two hypotheses from the maximum VMC (Velocity Made Good on Course) strategy. This basic sailing strategy involves sailboats traveling in the direction maximizing their VMC, which is defined as the velocity component parallel to the goal (Fig. 3A) (36). Under crosswind conditions, the optimal VMC direction aligns approximately with the goal direction (Fig. 3B). Conversely, when the goal is downwind (leeward) or upwind (windward), two high VMC directions deviating from the goal become apparent (Fig. 3B). This orientation also affects large-scale movement (Fig. 3C). Under crosswind conditions, the VMC-maximizing route tends to be a straight line. When the goal is downwind or upwind, sailors need to periodically change travel directions, a maneuver known as ’tacking’—leading to zigzag track patterns.

**Fig. 3.**
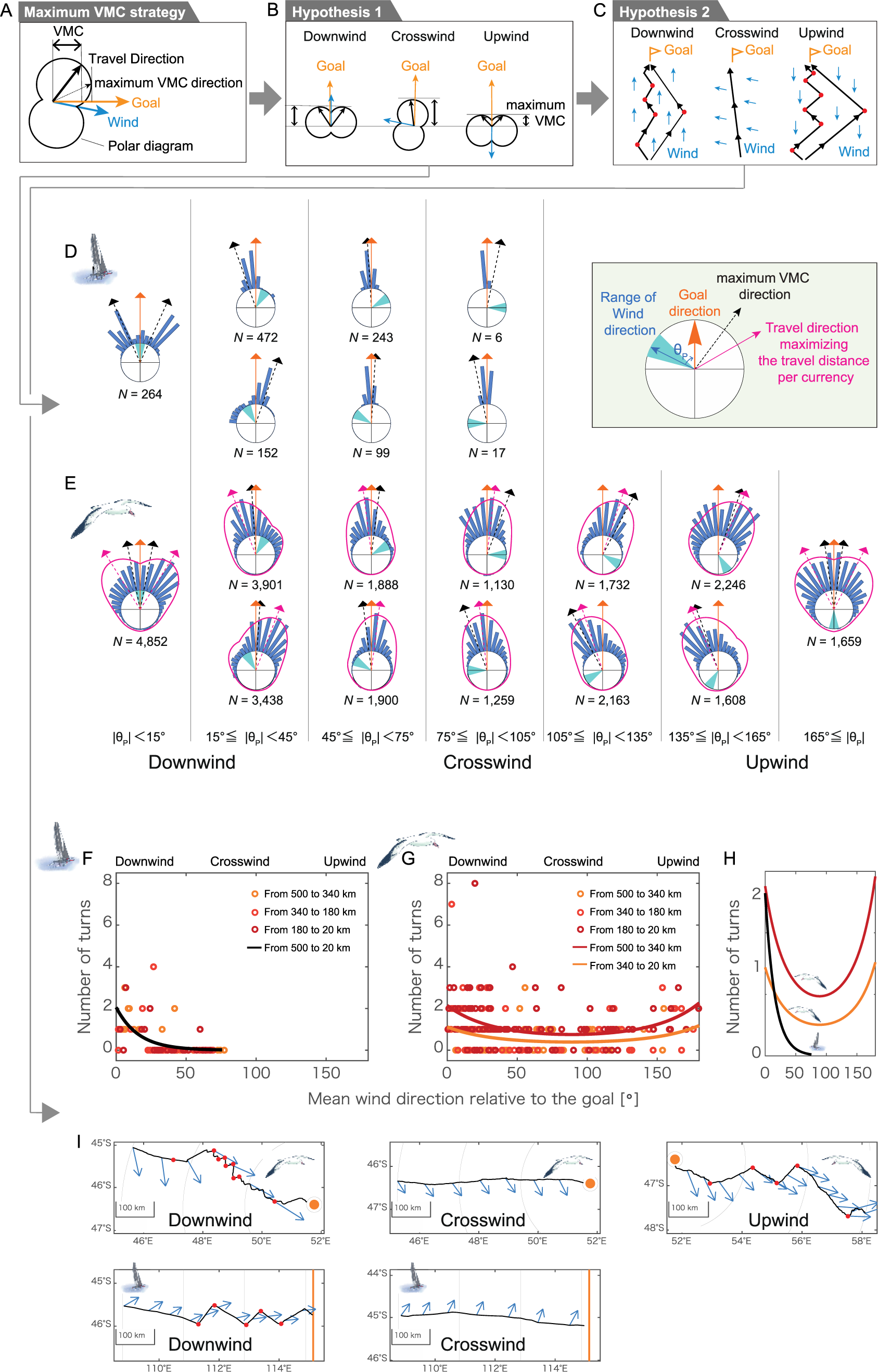
Prediction of albatross movement based on time-minimizing orientation strategy by sailors and test in track data. (A) Definition of velocity made course (VMC). (B) Travel directions that maximize VMC for each wind condition. Blue arrows indicate wind direction, orange arrows indicate the goal direction, and black arrows indicate travel directions that maximize the VMC. (C) Macroscale travel patterns predicted from the maximum VMC strategy. Red points represent turning points. (D–E) Histograms of the travel direction of sailboats (D) and albatrosses (E) relative to the goal direction. Each row indicates the different wind directions relative to the goal direction. The histograms are generated from the track data in Fig. 1 (within 500 km from the finish line or nesting island). The orange arrows indicate the goal direction. The cyan fans indicate the range of wind directions. The black arrows indicate the maximum VMC direction. For the albatrosses, the travel directions that maximize the travel distance along the goal per cost are shown with pink arrows. In addition, the fitted probability distribution of our stochastic model (see **METHODS** for detail) is shown by pink lines. (F–G) The number of turns in response to wind direction relative to the goal for sailboats (F) and albatrosses (G). (H) The fitted line of data in (F) and (G). Albatrosses evidently made more turns than sailboats. (I) Example of tracks (black solid lines) of albatrosses (upper row) and sailboats (lower row) within 500 km from their goals. The orange dots in the upper row represent the nesting island of birds. The orange line in the lower row shows the finish line (one of the middle checkpoints of the race corresponding to the longitude of Cape Leeuwin). The blue arrows represent the wind direction on the track. The red points represent the identified turning points. When the goal was located downwind and upwind (first and third column), more turns occurred compared to the crosswind condition (the second column).

To summarize our hypotheses, when the goal is located downwind or upwind, both yacht racers and albatrosses using a maximum VMC strategy are expected to deviate their travel direction from the goal (hypothesis 1) and increase the number of turns (hypothesis 2), compared to under crosswind conditions.

### Qualitatively similar movement patterns between albatrosses and sailboats

To test these hypotheses, we analyzed the tracks of sailboats and albatrosses. The Vendée Globe course begins and ends in Les Sables-d’Olonne, France, requiring a circumnavigation of the globe (Fig. 1E). En route, racers must cross several checkpoints, one of which is the 115°08’09’’E longitude line of Cape Leeuwin (Fig. 1F). For the purposes of this paper, we will refer to this line as the ‘finish line’, even though the actual race ends in Les Sables-d’Olonne. We used the tracks of sailboats within 500 km from the finish line. For albatrosses, we used the portion of their foraging trips homing to and within 500 km from the nesting island (Fig. 1B).

The data supported the first hypothesis. Figure 3D and E show histograms of the travel direction with different wind directions relative to the goal. These travel directions are every 2 min (*N* = 27,776) for albatrosses and every 30 min (*N* = 1,253) for sailboats. The peaks of the frequency distribution of travel direction for both sailboats and albatrosses deviated from the goal when it was located downwind; for albatrosses, this deviation also occurred when the goal was upwind.

The data also supported the second hypothesis. We defined a meander with >10 km width as one turn. Then, we counted the number of turns for three sections according to the distance from the goal (500–340 km, 340–180 km, and 180–20 km). The average wind direction of each section was also determined. We obtained 226 sections from 95 tracks for albatrosses and 84 sections from 28 tracks for sailboats (See **METHODS** for details). The number of turns increased significantly with the downwind goal for both sailboats and albatrosses and with the upwind goal for albatrosses (Fig. 3F–H and Table S1–S2). Examples of zigzag patterned tracks in downwind and upwind goals, as well as those of straight patterned tracks in crosswind goals, are shown in Fig. 3I. All tracks are shown in Fig. S2–S7.

### Quantitative difference between movement patterns of albatrosses and sailboats

While our analysis showed qualitative similarities between the movement patterns of albatrosses and yacht racers, it also implied some quantitative differences that were further examined as follows.

#### (i) Deviation from time minimization as primary travel cost

The observed travel direction of albatrosses slightly deviated from the time-minimizing flight directions (black arrows in Fig. 3E). This deviation implies that, for homing albatrosses, the time may not be the exact travel cost, a variable that animals attempt to minimize during their travel. For example, energy may be the true travel cost. Recall that when we employed the time-minimizing hypothesis for albatrosses, we assumed their travel time to be a good proxy for energy expenditure. Thus, we have implicitly assumed that the time-minimizing strategy and energy-minimizing strategy are identical. However, given that the heart rate—an indicator of energy expenditure—in wandering albatrosses increases with headwind (21), the travel direction minimizing energy expenditure could diverge from the time-minimizing direction. This difference may explain the observed deviation.

To investigate the characteristics of the travel cost for albatross, we developed a stochastic movement model (see **METHODS**). Our model postulate a bird moves with a higher probability in the direction yielding a greater travel distance per cost along the goal direction. This probability is governed by the ‘cost function’ which illustrates how the cost expenditure per time varies with the angular difference between the bird’s travel direction and wind direction. By testing various cost functions, we can gain insights into the characteristics of cost factors that well explain the observed travel direction. Here, we tested three cost functions that represent potential costs: constant (implied cost: time), linear (implied cost: energy), and quadratic (implied cost: unknown). See *Formulation of cost functions* in **METHODS** for detail. Models assuming each function were applied to the track direction data, and the quadratic function model was selected based on the Bayesian information criterion (BIC). The estimated cost function increased with headwind and tailwind (Fig. 4A and Table S3). This outcome suggests that the travel cost is not solely time or even energy and may incorporate other factors.

**Fig. 4.**
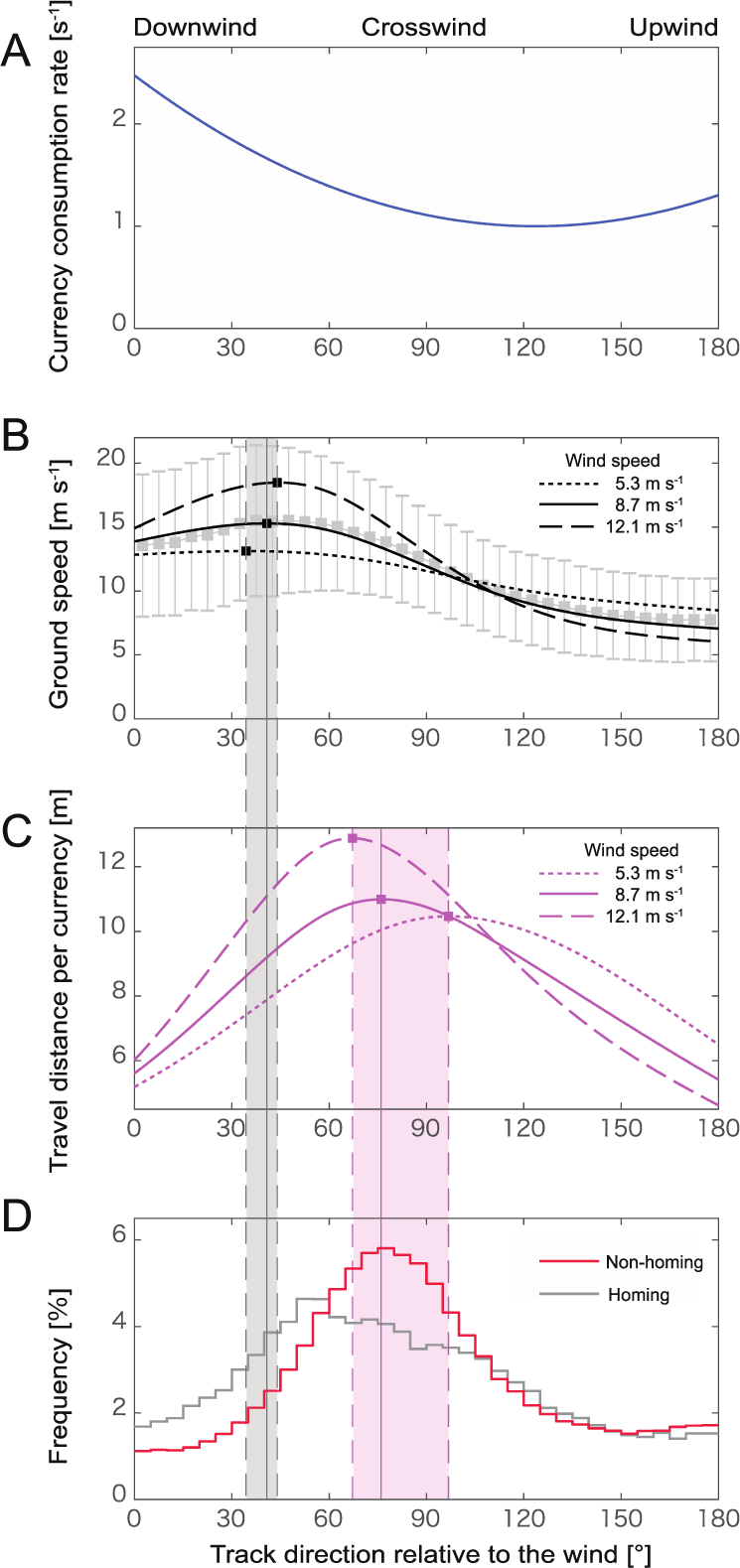
Travel direction of albatrosses deviates from the speed maximizing direction. (A) Estimated cost consumption rate function, *C*(*θ*_*G*_) = 1 + 0.314(|*θ*_*G*_ | − 2.18)^2^. The horizontal axis represents the travel direction relative to the wind. (B) The mean and standard deviation values of the ground speed of wandering albatrosses for all tracks are shown for each 5° of travel direction relative to the wind. The black lines correspond to the polar diagram in Fig. 2. The solid black line represents a wind speed of 8.7 m s^−1^ (the mean wind speed experienced by the albatross), the dotted line of 5.3 m s^−1^, and the dashed line of 12.1 m s^−1^ (mean wind speed ± standard deviation). The black squares represent the travel directions achieving the maximum ground speed for each wind speed. (C) The distance an albatross can travel per cost at a wind speed of 8.7 m s^−1^ (solid line), 5.3 m s^−1^ (dotted line), and 12.1 m s^−1^ (dashed line). These lines are obtained by dividing the polar diagrams in (B) with the cost function in (A). The maximum value is obtained when the travel direction of the bird to the wind is 83°. (D) Histogram of the travel direction relative to the wind for wandering albatrosses in the non-homing (red, *N* = 374,969) and homing (right gray *N* = 47,839) phase. The grey zone on panel B-D, indicates range of the travel directions achieving the maximum ground speed at wind speed from 5.3 m s^−1^ to 12.1 m s^−1^. The pink zone on panel C-D, indicates range of the travel directions maximizing travel distance per the cost. The peak of the frequency distribution of the travel direction to the wind is out of the gray zone, but well within the pink zone.

In summary, while time serves as a reasonable proxy for travel cost given its efficacy in explaining the qualitative movement patterns of homing albatrosses, the quantitative analysis suggests the need for incorporating additional factors into the cost. Potential factors will be discussed in the **Discussion**.

#### (ii) Difference in frequency of large-scale turns

Quantitative differences in the turning frequency during large-scale movement were also observed (Fig. 3H). First, the turning frequency was higher in albatrosses than in sailboats. Second, the turning frequency for sailboats showed no correlation with the distance to the goal, whereas albatrosses exhibited a higher turning frequency when the goal was closer (340–20 km) compared to when it was farther away (500–340 km, see Table S1–2).

## Discussion

This study is the first to compare the track data of wandering albatrosses and sailboats. Our results reveal that both demonstrate similar orientation and routing when heading towards their goals, albeit with some quantitative differences.

### Qualitative findings: albatrosses show similar orientation as sailboats and ‘tacking’

Understanding how birds adjust their travel direction in response to the wind directions relative to their goal directions is fundamental to comprehending their movement strategies (11). However, the interrelation between travel direction, wind direction, and goal direction has rarely been explored in dynamic soaring birds. Although previous studies have reported that dynamic soaring birds prefer tail-to-side winds (8, 21, 26, 27, 29, 30), these studies primarily examined the relationship between only the wind direction and the bird’s travel direction. Our study found that, similar to sailboat racers, albatrosses flexibly adjust their travel direction based on the goal direction relative to the wind. They deviated from leeward and windward goals in small-scale movements (1–2 km) and switched direction more frequently when the goal is upwind or downwind in large-scale (>10 km) movements, with some tracks showing clear zigzag patterns. It should be noted that this large-scale zigzag completely differs from the well-known several-hundred-meter-scale zigzag pattern of albatrosses (40–42), which stems from the S-shape track of one cycle of dynamic soaring.

To the best of our knowledge, this is the first empirical evidence that albatrosses adjust their direction of movement in response to wind direction similar to sailboat racers. Two studies (28, 41) have reported that tracks of albatross proceeding upwind exhibit zigzags at 100 m to 1 km scales, and this phenomenon was discussed using yacht tacking as an analogy (23, 41). However, these discussions were based on visually selected data from the complete datasets, with the total combined length of the selected tracks approximating 20 km. Additionally, a recent study (30) reported ’zigzag flights’ during the foraging trips of blue petrels. This zigzag flight, observed in the non-homing phase, consists of approximately 180° turns, resulting in the birds staying within a restricted area. The zigzag pattern was considered a strategy for increasing prey search efficiency in situations without a specific goal (30), because flying crosswind can enhance the travel distance of birds per unit time or energy (23, 30, 41). Thus, this zigzag flight stems solely from a preference for crosswinds and is different from ’tacking,’ which arises from a tension between the preference for crosswinds and the necessity to move towards a specific goal. Consequently, to confirm ’tacking’ in dynamic soaring birds, merely identifying zigzag patterns in their flight tracks is not enough; it is crucial to quantify how the occurrence of zigzag patterns changes depending on wind and goal directions during the homing phase (i.e., whether the wind is blowing headwind, downwind, or crosswind to the goal). By considering these factors, we presented the first experimental evidence of tacking behavior in albatrosses.

### Quantitative findings

#### (i) Albatrosses make more turns than sailboats

Our study revealed that albatrosses turned more frequently than sailboats. This could be due to differing turning costs for sailboats and albatrosses. Turning in sailboats requires adjusting the sail’s direction, a maneuver potentially more time and energy-consuming than turns made by albatrosses. Consequently, yacht racers likely adopt strategies to reduce the number of turns. We also found an increase in the turning frequency as albatrosses neared their goals, a trend absent in sailboats. This might be due to the difference in their navigation goals: for sailboats, the goal is typically a line, providing some flexibility, whereas for albatrosses, it is a specific point.

#### (ii) Albatrosses do not maximize their speed

Our findings suggest that the travel cost for albatrosses during their homing phase cannot be attributed solely to time or energy. This challenges the prevailing understanding that dynamic soaring birds, such as albatrosses, favor crosswinds to optimize their travel speed (8, 21). The prey of albatrosses is widely and randomly distributed over the sea. Therefore, during their foraging trips (excluding the homing phase), the priority should be to maximize the encounter rate with prey. This implies that albatrosses are expected to choose a direction that optimizes travel distance per unit cost during the non-homing phase. A previous study demonstrated that wandering albatrosses prefer the speed-maximizing direction during foraging trips (21), suggesting that their travel cost is time (or energy). Comparable findings have been reported for Gadfly petrels, another dynamic soaring species (8).

However, these studies analyzed the entire trip and thus did not differentiate between the non-homing and homing phases. Our results suggested that time was not the sole cost during the ’homing’ phase for wandering albatrosses. To reconcile our findings with those of previous studies, we propose two possible hypotheses. The first hypothesis is that time is the travel cost during the non-homing phase (based on previous studies) but not during the homing phase (based on our results). The second hypothesis is that the travel cost cannot be attributed solely to time even during the non-homing phase. Although the second hypothesis contradicts the results of previous studies, it is worth noting that the tracking data used in those studies had a coarse temporal resolution of 1-to 2-hour sampling intervals (8, 21). This could potentially obscure the detection of small-scale movement patterns.

We calculated the travel direction of albatrosses during the non-homing phase from our data, recorded at 2-minute intervals (Fig. 4D). Unlike coarser resolution tracks used in previous studies, our data indicated that the peak of travel direction distribution deviated from the speed-maximizing direction (black lines in Fig. 4D). However, it closely aligned with the direction that maximizes travel distance per the cost which was estimated from the homing segments of the tracks (pink lines in Fig. 4D). Therefore, these results support the second hypothesis. Our data suggest that albatrosses incur the same cost during both the non-homing and homing phases, implying the possible contribution of factors other than time or energy to the travel cost throughout their foraging trip.

### Speed or robustness: Which will be the priority for the dynamic soaring birds in fluctuating winds?

What then constitutes the additional cost? While our current results cannot identify it definitively, one potential contributor could be the risk of costly flapping flights due to the unpredictability of wind conditions. Soaring birds are spared from flapping flight by exploiting the energy from wind. However, dynamic and unpredictable wind conditions may necessitate birds to engage in costly flapping flights. Hence, for soaring flight, not only efficiency (i.e., time or energy minimization in predictable wind) but also the robustness to stochastic changes in the winds should be key factors, whereby birds try to minimize the duration of flapping flight. This is particularly important for larger species such as the wandering albatross, which rarely uses flapping flight (17).

This risk-aversion strategy is already known in thermal soaring birds (43–45). Thermal soaring is a flight style in which birds repeatedly ascend with updrafts (convection currents) and then glide. If birds are aware of the distance and updraft speed of successive thermals, the theory suggests an ’optimal speed’ to maximize their horizontal travel speed (46, 47). However, in practice, larger bird species tend to employ a slower airspeed than the optimal speed (43). The slower airspeed allows them to cover longer distances with a smaller loss in altitude (in contrast to that optimal speed enables longer distance with shorter time). This tactic mitigates the risk of the birds losing height before finding the next stochastically distributed thermal, thus avoiding the risk of being forced to conduct costly flapping flights (43).

Our results may suggest that dynamic soaring species may also employ a risk-aversion strategy. The theory of dynamic soaring often assumes wind gradients to be temporally invariant (20, 48, 49). However, real wind gradients are turbulent and exhibit spatiotemporal fluctuations (50). Moreover, dynamic soaring birds fly near the sea surface, where the wind gradient becomes quite complicated due to the interplay between wind and waves (51). This uncertainty in wind conditions may drive albatrosses to prioritize the efficiency of harvesting mechanical energy from wind gradients over flight speed. In dynamic soaring, a trade-off exists between speed and the mechanical energy harvesting from the wind (9): the equation of motion shows that the kinematic energy flow from wind gradient to the bird increases when the bird aligns its flight direction upwind during ascent and downwind during descent (9, 19, 20). Therefore, compared to a strategy that prioritizes energy harvesting, a strategy that maximizes speed may be more prone to unexpected wind changes and carry a higher risk exhausting mechanical energy needed to stay aloft, potentially leading to costly flapping flights. To avoid this risk, wandering albatrosses may favor travel directions that prioritize energy-harvesting efficiency, even if it results in slower speeds. Future work could focus on testing whether dynamic soaring birds prioritize energy-harvesting efficiency over travel speed. This would be an intriguing prospect, but it is technically challenging due to the need for the simultaneous recording of detailed tracking, body posture, and air speed of the bird at high resolution.

### Navigation strategies of yacht racers provide a new perspective on seabird navigation

Pelagic seabirds are unique in the animal kingdom, with their extensive track data offering invaluable opportunities to explore wildlife movement in response to the surrounding environment (25, 52–54). This study demonstrates that insights from sailboat racers and engineers can aid in decoding these extensive data to reveal fundamental patterns of pelagic seabird locomotion, laying the groundwork for further exploration. While our study simplistically assumed that albatrosses choose their travel direction based solely on local wind, birds may perceive or predict larger-scale wind conditions and adjust their travel direction accordingly (8, 21, 30). Autonomous sailing algorithms could be useful to further investigate these intricate movement strategies (55, 56). Such algorithms determine the optimal travel direction by incorporating global-scale wind predictions (55, 56). Applying these man-made movement strategies as testable hypotheses to pelagic seabirds’ movement could help uncover the travel costs and cognitive capabilities of these species, thereby offering interesting directions for future investigation.

## METHODS

### Track data

We used the track data of wandering albatrosses from 2003–2005 and 2016–2019 collected in previous studies (33, 34, 57) (Fig. 1A–D). All data were obtained from breeding individuals on Possession Island (46°25’S. 51°45’E). Loggers were attached on the back feathers with TESA tape during a shift change with their partner. Loggers were retrieved when they returned to the nest after foraging. Loggers weighed 60-75 g in the 2016-2019 studies (34, 57); in the 2003-2005 study (33), in addition to the 45 g GPS logger, birds swallowed a 20 g stomach temperature tablet that transmitted stomach temperature to a recorder, and the 25 g recorder was attached to their backs, for a total device mass of 90 g. In all studies, the total mass of the equipment was less than 3%. See the each study for more detailed information (33, 34, 57)

Portions of tracks within 20 km of the Island were excluded from the analysis to avoid the influence of land. Trips of incomplete recordings that stopped before the nest was reached were excluded from the analysis. In total, we obtained data for 149 foraging trips. An iterative forward/backward averaging filter was applied to each track to exclude unrealistic points with speeds of more than 100 km h^−1^. Furthermore, since sampling intervals differed among the data (10 s to 2 min), the data were resampled every 2 min.

Additionally, data from the Vendée Globe, a long-distance yacht race held in 2020, was used for sailboats (Fig. 1E–H). Of the participating boats, we used data from 28 boats that passed through the middle checkpoint (longitude 115°08’09’’E line). For the analysis, we used the route below a latitude of 30 degrees north. The positions of all the boats were recorded every 30 min.

### Defining the homing phase of wandering albatross

We identified the homing start points for each albatross track. The portion of tracks after the homing start point was defined as the homing phase, while that prior to the start point was defined as the non-homing phase. We employed a backward path analysis to determine the homing start point (58, 59). In this method, by starting at the goal location and moving backward along the path, the backward beeline distance (*BD*) and the backward path length (*BL*) were calculated for each point. The point at which *BD* stopped increasing linearly with respect to *BL* was defined as a homing starting point. Although previous studies visually determined the point of change from linear to non-linear, in this study, we defined the homing starting point using the following procedure to ensure reproducibility. First, we calculated the *BD* and *BL* for each point. The *BL* and *BD* data set was then resampled to record BL for every 10 km. We then applied a 50 km moving average to the *BL*. Subsequently, *BD* was plotted against *BL*, and the peak points of *BD* were detected. Among these peaks, point *P* with the smallest *BL* was determined. In the *BD* vs. *BL* plot, a linear and broken line with one change point was respectively fitted to the region where *BL* ≤ *P*, and the BIC were calculated for the two. If the broken line showed a lower BIC than the linear line, and the slope of the line that was further away from the nest was less than half of the slope of the line that was closer to the nest, the position corresponding to the change point of the broken line was determined as the homing start point. In all other cases, the position corresponding to point *P* was determined to be the homing start point.

### Calculation of ground velocity vector, wind vector, and goal direction

For each data point, we calculated (i) the ground velocity vector (travel direction and speed to the ground) and (ii) wind direction and wind speed. In addition, for data points in the homing phase and within 500 km of the goal, we also calculated (iii) the direction of the goal as follows.

#### (i) Ground velocity (ground speed and travel direction)

The ground velocity was calculated for each position by dividing the vector connecting two consecutive data points by the elapsed time. Since there were some instances of recording deficit, the ground velocity was not calculated for data points corresponding to these deficits; i.e., the ground velocity vectors were calculated only when there was a pre-resampling data point within 2 min before and after the two consecutive post-resampling data points. The track data included data when the albatross was staying at the sea surface. Therefore, albatross data points with a speed of less than 2 m s^−1^ were regarded as data points during their stay at the sea surface and were excluded from the analysis.

#### (ii) Wind direction and speed

For the wind direction and speed data, we used ERA5 ECMWF, hourly on a 0.25° grid (https://cds.climate.copernicus.eu/cdsapp#!/dataset/reanalysis-era5-single-levels?tab=overview). For each track data point, the estimates of wind direction and speed predictions at the closest point in time and distance were used.

#### (iii) Goal direction

The direction of the goal from the bird or sailboat was calculated for each data point for data within 500 km from the goal in the homing part of albatrosses and within 500 km from the finish line of the sailboat. For the albatrosses, the goal direction was calculated from GPS observation points, setting Possession Island as the goal, and for the sailboats, the goal direction was set to due east.

### Identification of polar diagram using GAM

Based on the ground velocity vector and the wind vector obtained in the previous section, the polar diagrams of the albatrosses and sailboats were determined using a generalized additive model (GAM), a non-parametric smoother method (60). The ground speed (*V*) was used as the response variable, and the absolute value of the direction of movement relative to the wind (*θ*_*G*_: difference between the travel direction and the wind direction, see Fig. S8) and the wind speed (*w*) were the explanatory variables. The calculations were performed in R v3.6.3 with the ‘gam’ function of the ‘mgcv’ package. We employed the ‘te()’ function setting the tensor product smooths for the model formula (60). From these, the ground speed was obtained as a function of the direction of movement and wind speed relative to the wind, i.e., *V*(*θ*_*G*_, *w*), and the polar diagram was obtained by displaying this function in polar coordinates.

### Counting turns in tracks

To quantify the number of large-scale turns (>10 km wide), we applied the following procedure to portions of track data from sailboats and albatrosses within 500 km and >20 km from their goals.

1. For distances less than *x* = 500, 420, 340, 260, or 180 km, at which the albatrosses started homing.

1-1 : A point of track *x* km from the goal was selected as well as a point at (*x* - 160) km from the goal, and a beeline was drawn by connecting these two points (Fig. S1A).
1-2 : A time series of the **PDB** (Perpendicular Distance of the position of the traveler from the Beeline) was computed for points from *x* to (x - 160) km from the goal point (Fig. S1B). The PDB was defined as positive when the traveler was on the left side of the beeline and negative when the traveler was on the right side. When the track is straight, the PDB does not show a clear peak. In contrast, when the track is zigzag, PDB shows a distinct peak. The number of peaks in the PDB time series corresponds to the number of turns, and the degree of change represents the width of the zigzag pattern. Therefore, in this study, we detected the large-scale turns from the PDB. Specifically, we performed the following two procedures.
2. For each PDB time series obtained in (1), the peaks with a prominence of 10 km or more were calculated at the bottom (blue points in Fig. S1C–D) and top (green points in Fig. S1C–D). The prominence of a peak is an index to quantify how much a peak stands out from the surrounding baseline of the signal, and is defined as the vertical distance between the peak and its lowest contour line. The ‘findpeaks’ function in MATLAB v2019a was used to detect the peaks.
3. For the detected peaks, if there were consecutive peaks with the same side (top-top or bottom-bottom), the one with the highest absolute value of the peak was selected (Fig. S1E–F). The points selected through these procedures were defined as ‘turns’ (red points in Fig. S1E–F).

We tested the effect of wind direction and distance from the goal on the occurrence of these turns. First, the number of turns per 160 km in three sections (500–340 km, 340– 180 km, and 180–20 km from the goal) were counted. Then, we determined the mean goal direction to the wind; defined as the absolute difference between the mean wind direction and the mean goal direction experienced by the bird or sailboat every 160 km. We fitted ‘glm’ by setting the mean goal direction to the wind (from 0 to π rad) and the section of the movement (three categorical variables 500–340 km, 340–180 km, and 180–20 km) as the explanatory variables, and the number of turns in the 160 km section as the response variable. The log link function and the Poisson distribution were employed. To capture the feature that the number of turns increased with a tailwind and upwind, the square of the mean goal direction to the wind was added to the linear predictor in addition to the average direction of the goal to the wind and the section of the movement.

### Model to estimate travel cost from track data

We have developed a stochastic movement model to determine the ’travel cost’ that best explains albatross movement data. Our model considers bird flight in a two-dimensional space. Here, we denote wind speed as *w*, the angular difference between the bird’s ground velocity and the wind velocity as *θ*_*G*_, and the angular difference between the goal direction and the wind velocity as *θ*_*P*_ (Fig. S8). Our model is based on the premise that a bird is more likely to move in a direction that closely aligns with maximizing the ’distance traveled in the goal direction per unit of cost spent’. This likelihood is determined by a cost function that quantifies the rate of cost expenditure depending on *θ*_*G*_. The aim of our model is to estimate the form of this cost function using empirical data, thereby gaining insights into the nature of the travel cost for albatrosses. Firstly, we present the model formulation to describe the bird’s travel direction (Step 1), then we discuss potential options for the cost function (Step 2), and finally, we estimate the cost function by fitting the model to the data (Step 3).

#### [Step 1] formulation of the model

We define the cost function *C*(*θ*_*G*_) as the cost consumed per unit time during a bird’s flight in the direction of *θ*_*G*_. We assume the cost function only depends on *θ*_*G*_ for simplicity. The specific form of the cost function will be discussed in subsequent steps.

We then formulate the ’distance traveled in the goal direction per unit of cost spent’ using the cost function. For this purpose, we used the concept of Isotropic Energy Polygons (IEPs) proposed in a previous study (16). IEPs graphically represent the distance an animal can travel per unit energy spent. In our case, i.e., two dimensional bird flight, the IEP is a polar plot of *L*_*E*_(*θ*_*G*_, *w*) with respect to *θ*_*G*_, where the function *L*_*E*_(*θ*_*G*_, *w*) represent the distance covered by a bird per unit energy spent under given travel direction *θ*_*G*_ and wind speed *w*. Following this idea of IEPs, we propose Isotropic Cost Polygons (ICPs), a generalized version of IEPs that is characterized by the distance a bird can travel per unit of ‘cost’ spent. Hence, an ICP is a polar plot of *L*_*C*_(*θ*_*G*_, *w*) which represents the distance covered per unit cost. Using ICPs (see Fig. S8), the ’distance traveled in the goal direction per unit of cost spent’, denoted as *F*(*θ*_*G*_|*θ*_*P*_, *w*), is given by:

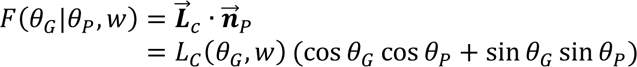

Here, 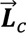 denotes a vector that connects the origin to a point on the ICP when the direction of movement relative to the wind is *θ*_*G*_, and 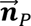 denotes a unit vector heading to the goal direction. The distance traveled per unit of cost, denoted as *L*_*C*_(*θ*_*G*_, *w*), is equal to the distance traveled per unit time (i.e., the ground speed *V*(*θ*_*G*_, *w*)) divided by the amount of cost consumed per unit time (i.e., the cost function *C*(*θ*_*G*_)):

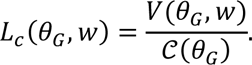

Thus, the following equation is derived:

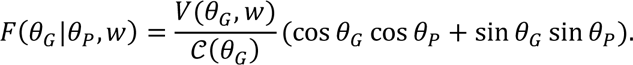

Finally, we assumed that probability distribution of *θ*_*G*_ was proportional to the exponent of *βF*(*θ*_*G*_|*θ*_*P*_, *w*), with *β* as a constant parameter:

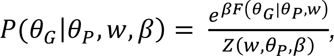

where *P*(*θ*_*G*_ |*θ*_*P*_, *w*, *β*) represents the probability distribution of *θ*_*G*_ given parameters *θ*_*P*_, *w*

and *β*. The term *Z*(*θ*_*P*_, *w*, *β*) represents the normalizing constant such that 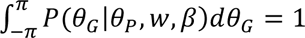, and thus is defined as 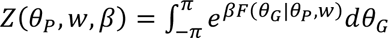.

From the above, we were able to model the birds’ direction of movement through the probability distribution *P*(*θ*_*G*_|*θ*_*P*_, *w*, *β*). This distribution is composed of the functions *V*(*θ*_*G*_, *w*) and *C*(*θ*_*G*_). The ground speed *V*(*θ*_*G*_, *w*) is already determined from the experimental data (see **Identification of polar diagram using GAM**). Thus, we need to define the specific form of the function *C*(*θ*_*G*_).

#### [Step 2] Formulation of cost functions

The cost function *C*(*θ*_*G*_) should ideally be simple while still capturing the characteristics of the assumed cost. To this end, we have formulated the cost function using polynomials up to the second-order of absolute value of *θ*_*G*_. We describe the assumed cost for each of the functional forms. These cost functions include some parameters, and we only allow parameters that keep the cost functions positive in the range of 0 ≤ |*θ*_*G*_| ≤ *π*.

For the **constant function**, where the cost is time,

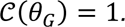

The consumed cost per time does not change depending on the travel direction. In this case, the travel cost is time.

For the **linear function**, where the cost is energy,

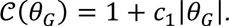

When *c*_1_ >0, the cost function increases when moving upwind, consistent with reports that heart rate, i.e., a good proxy of energy consumption rate, in albatrosses also increases under such conditions (21). This model, therefore, posits that energy constitutes the cost.

For the **quadratic function**. where cost is unknown,

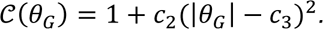

We assume that *c*_2_ >0. It reaches its minimum when moving in the direction of *c*_3_, and increases with any deviation from *c*_3_, whether upwind or downwind.

#### [Step 3] Stochastic model and calculation of the likelihood from the data

Our goal was to estimate the cost function that best explains natural bird movement data. For this purpose, the likelihood of the model on experimental data should be calculated. We denote the travel direction to the wind, wind speed, and goal direction to the wind at time *t* obtained from individual *i* as 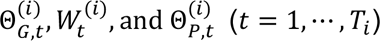, respectively. In this case, when the observation data is obtained from *n* individuals, the likelihood is given by

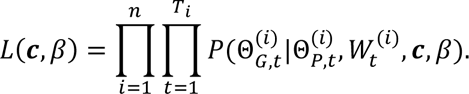

Where, *c* = {} for constant function, *c* = {*c*_1_} for linear function, and *c* = {*c*_1_, *c*_2_} for quadratic function. For each model that employed the cost functions described above, we computed the parameter ***c*** and *β* that maximizes this likelihood, and the BIC. Then we chose the cost function that best explained the data via model selection based on BIC. The ‘fminunc’ function in MATLAB 2019a was used. The normalization constants 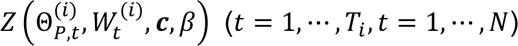 were computed numerically by Gaussian quadrature with 360 integration points. The estimated values of BIC and parameters for each model are shown in Table S3.

## Acknowledgements

The authors thank Chihiro Kinoshita for illustrating the albatross and sailboats in the figures. We thank Masanobu Katori for providing important information about sailing. We thank Vendée Globe for providing the track data of sailboats. We thank the fieldworkers involved in the study on Crozet, and in particular Julien Collet and Alexandre Corbeau. The field work was supported by IPEV (Program 109, PI HW). During the preparation of this work, we used ChatGPT to correct grammar. After interacting with ChatGPT, we meticulously reviewed and made necessary adjustments to the content. We take full responsibility for the content that has been published.

