## Supplementary for "Albatrosses employ orientation and routing strategies similar to yacht racers"

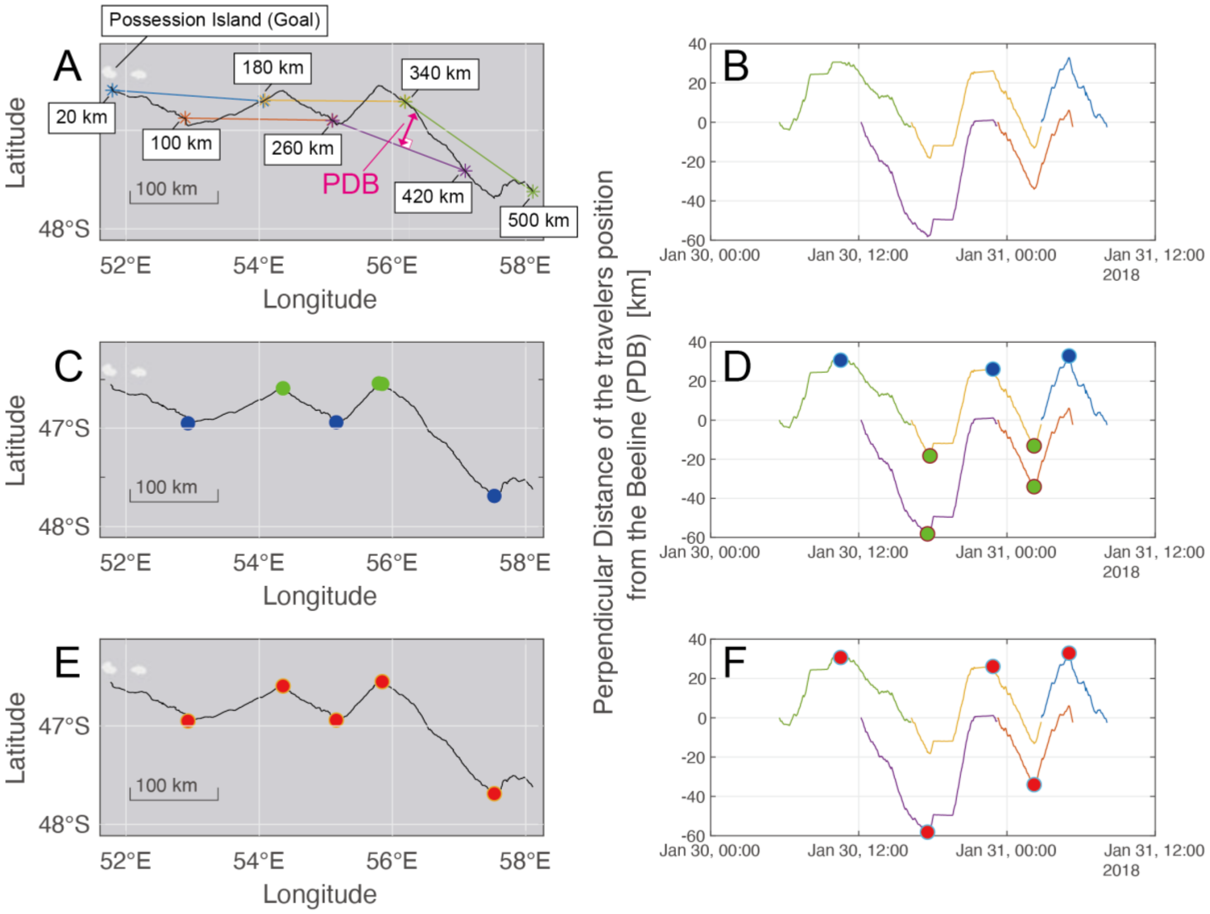


Fig. S1 Procedure for detecting turns. (A) Black line is the trajectory of the bird. The beelines connecting 500 km and 340 km from the goal on the path (green line), 420 km and 340 km (purple line), 340 km and 180 km (orange line), 260 km and 100 km (brown line), and 180 km and 20 km (blue line). The distance of the perpendicular line drawn from a location on the track to the beeline is PDB (pink line). (B) The time series of PDB was calculated for 500–340 km beeline (green), 420–260 km beeline (purple), 340–180 km beeline (orange), 260–100 km beeline (brown), and 180–20 km beeline (blue), respectively. (C) The locations that showed the peak of PDB are shown in (D). (E–F) locations defined as distinct turns.

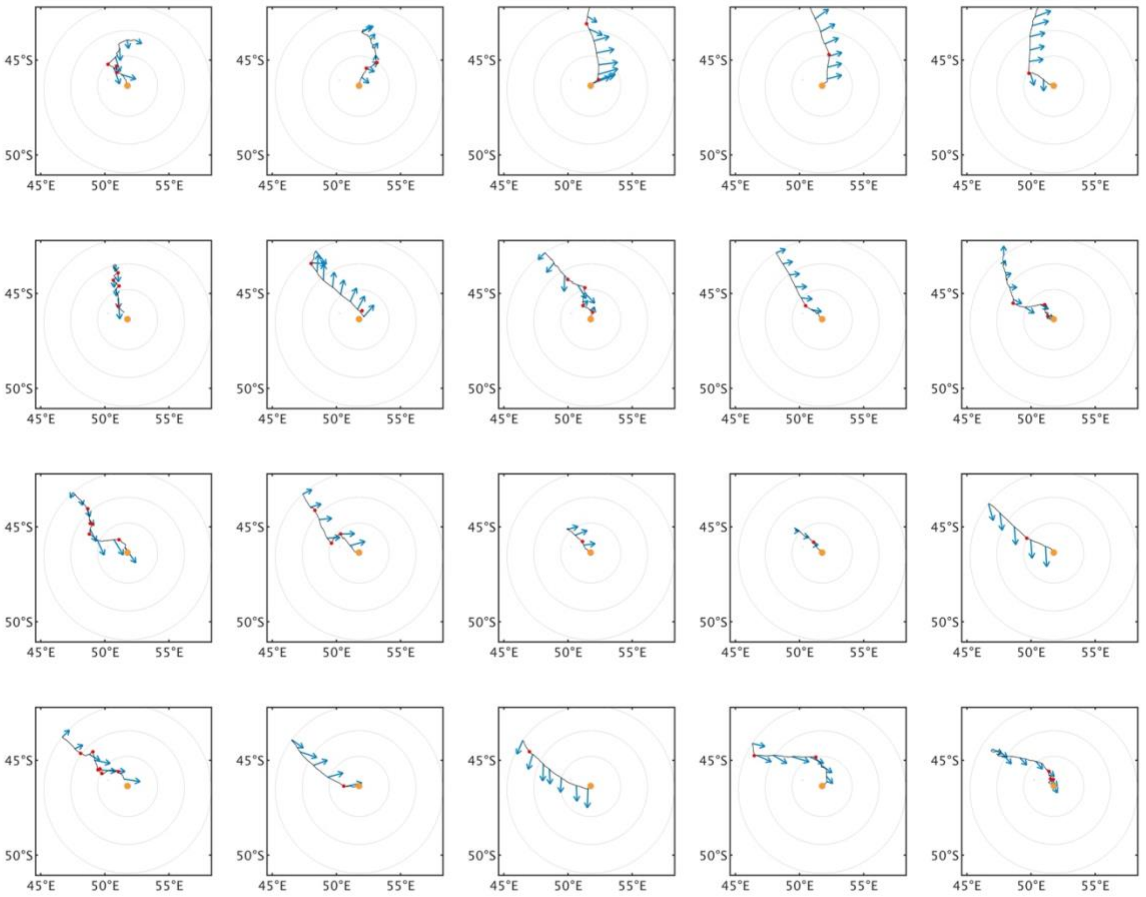


Fig. S2. Homing tracks of albatrosses that started their homing from a location more than 180 km away from the nesting island (orange point). The portion of homing tracks within 500 km of the island are shown. Gray circles are 20 km, 180 km, 340 km, and 500 km from the goal, respectively. Blue arrows are wind vectors at that time and location. Red dots represent locations identified as turn points. Tracks returning from the northwest side of the goal are shown.

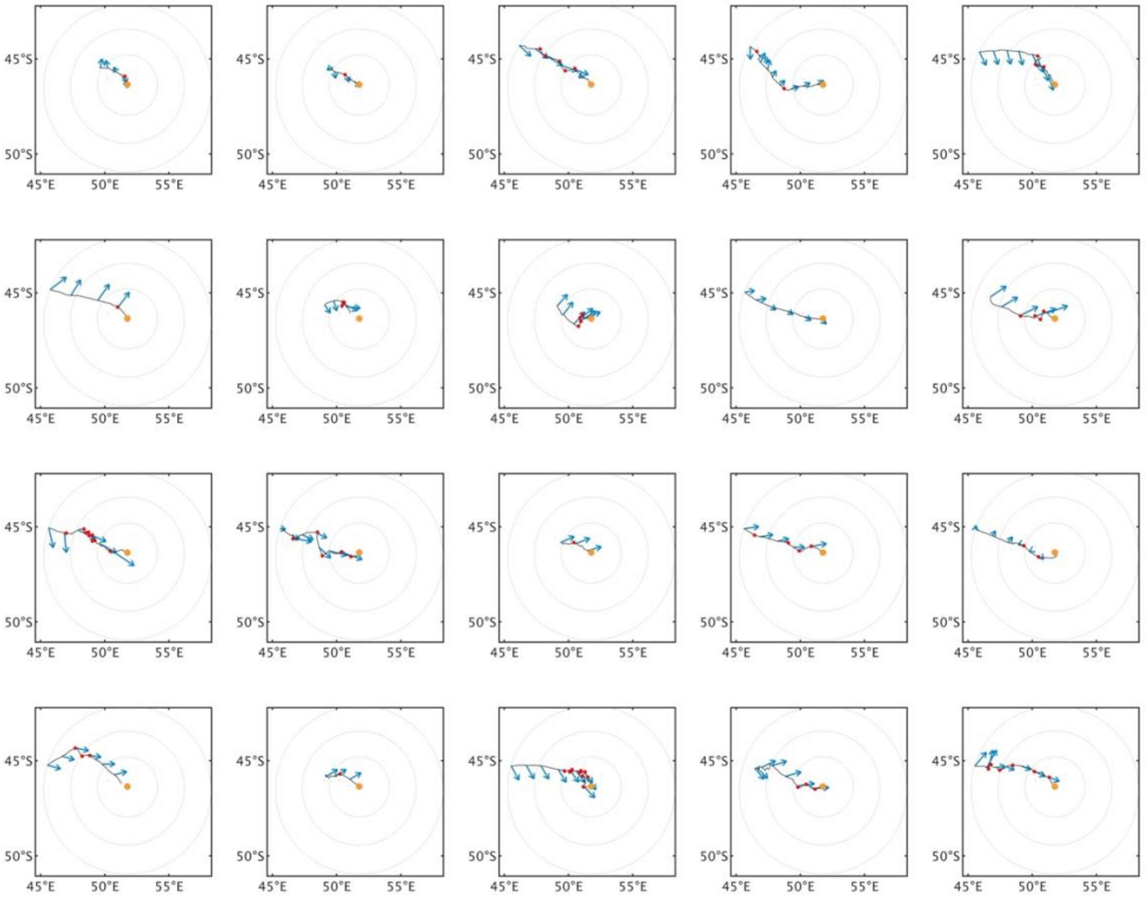


Fig. S3. Homing track of albatrosses, returning from the west side of the goal.

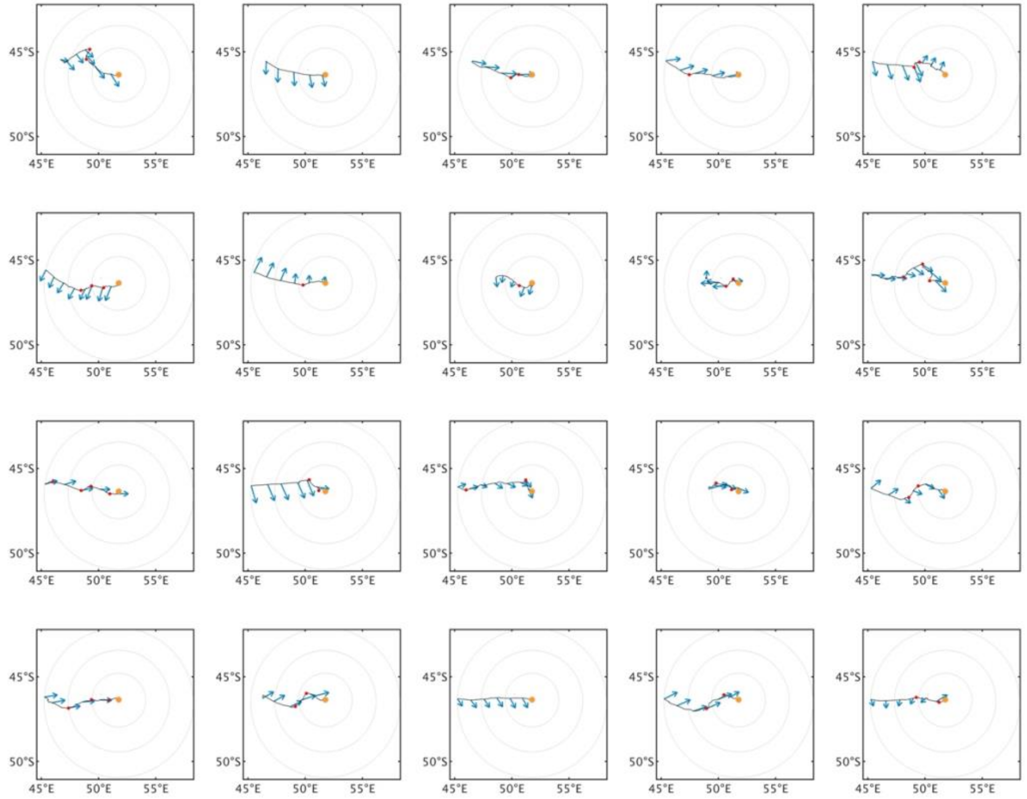


S4. Homing tracks of albatrosses, returning from the west side of the goal.

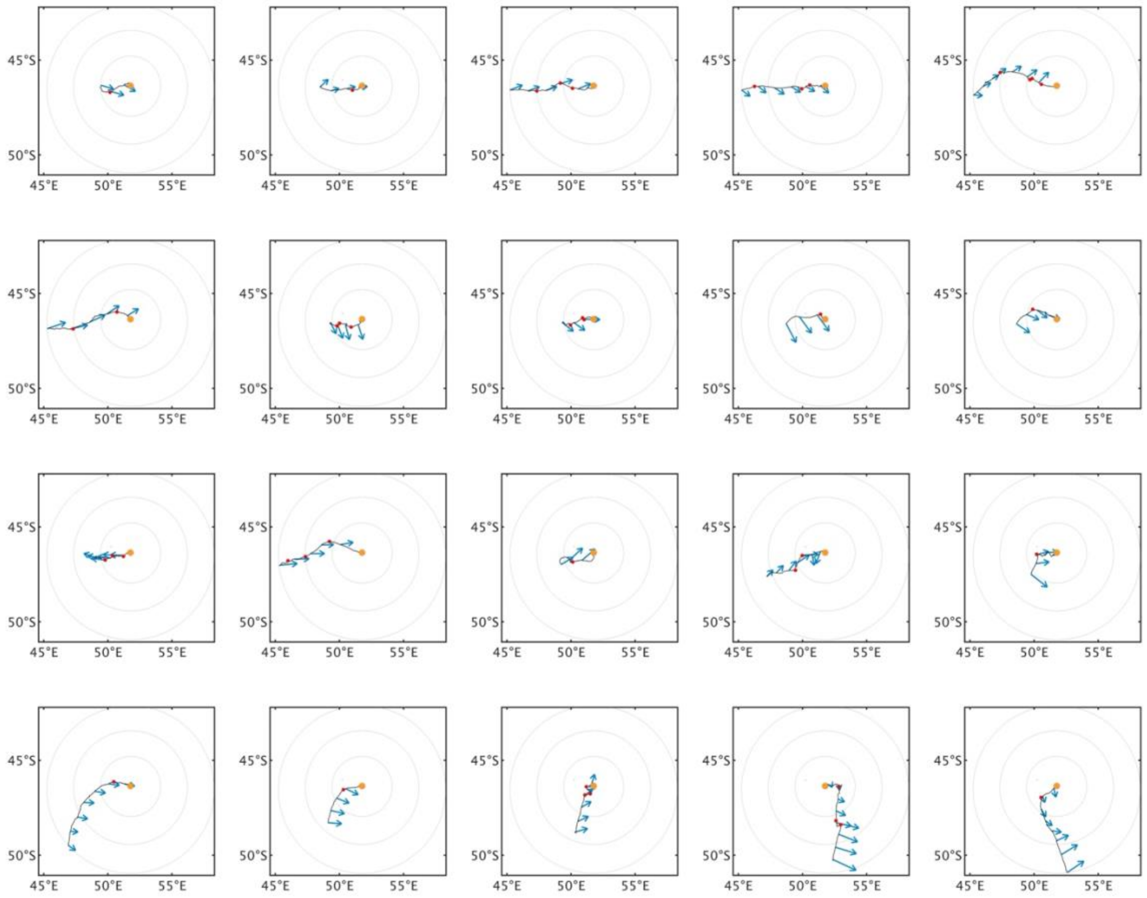


Fig. S5. Homing tracks of albatrosses, returning from the west south side of the goal.

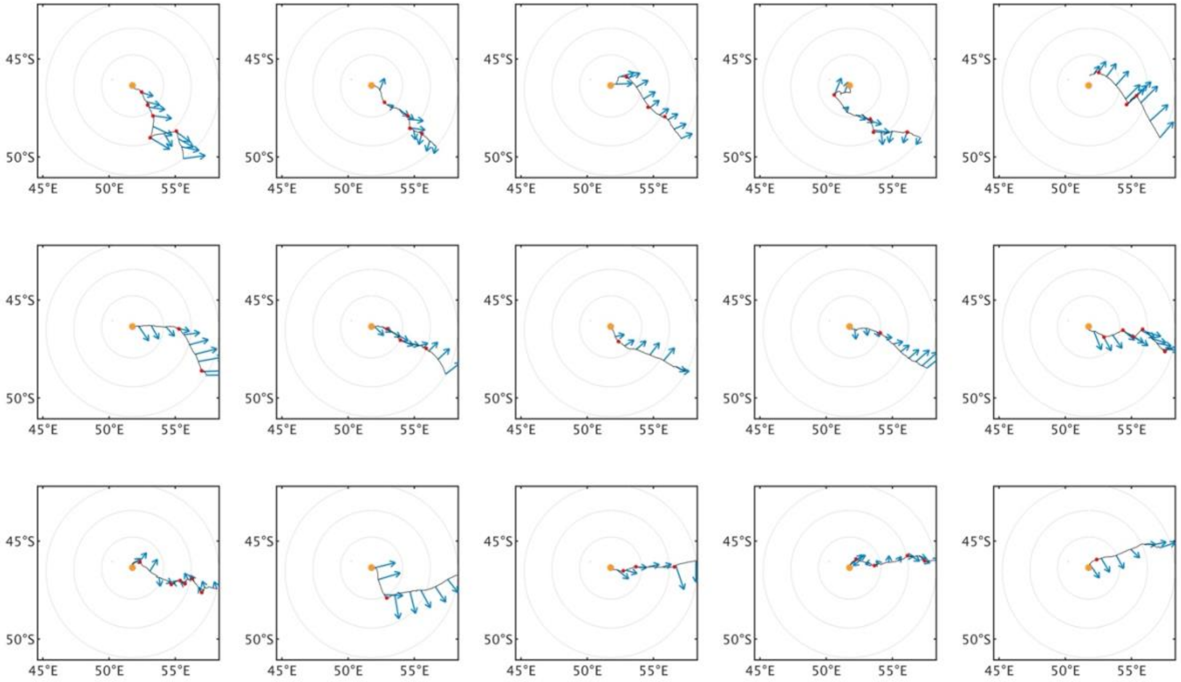


Fig. S6 Homing tracks of albatrosses, returning from the east side of the goal.

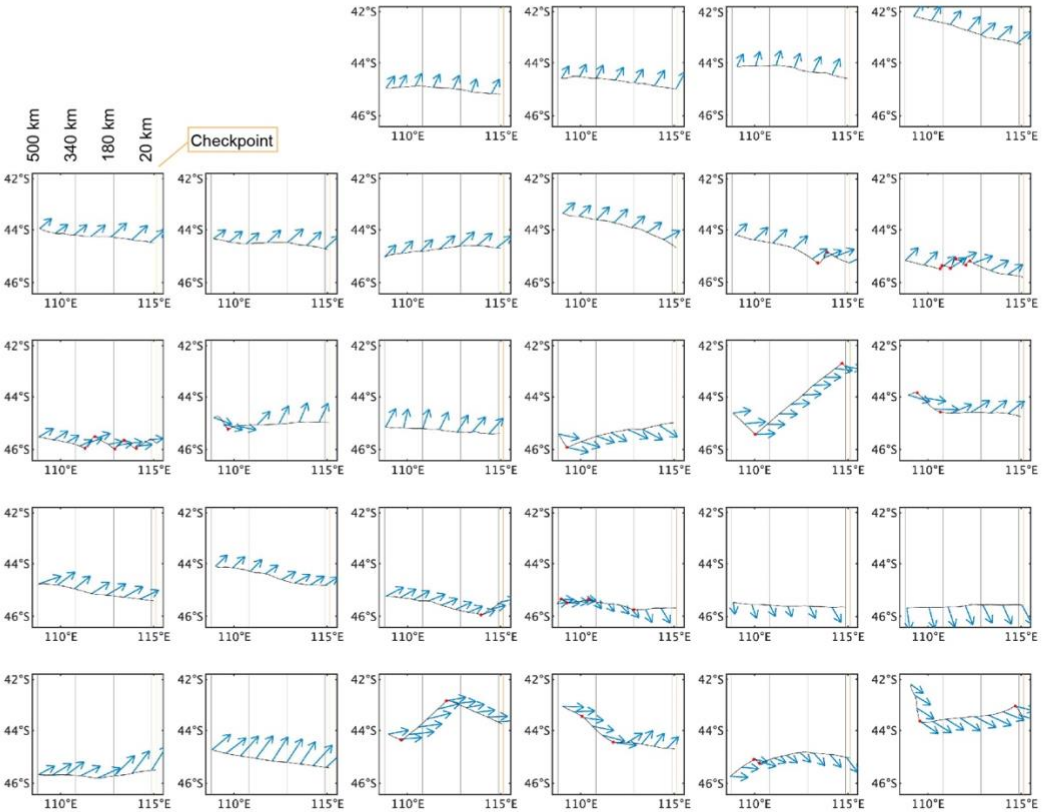


Fig. S7 Tracks of sailboats within 500 km of intermediate checkpoints. The orange line is the longitude line of the checkpoint. Blue arrows are wind vectors at that time and location. Red dots represent locations identified as turn points.

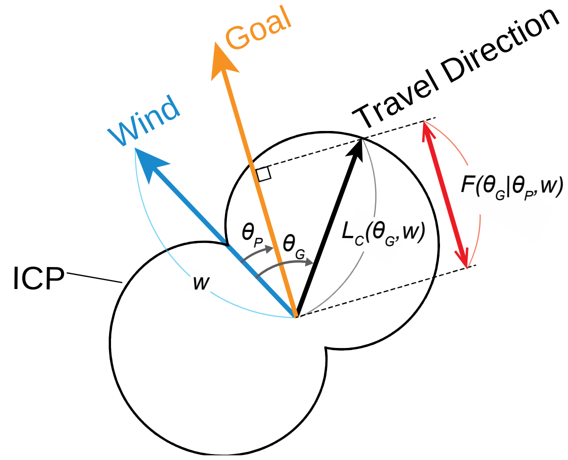


Fig. S8 Definition of terms and schematic diagram of ICP.

Table S1 Results of fitting glm models to data on the number of turns made by the albatrosses against the direction of the goal relative to the wind. The best model is shown in gray.

| Model | BIC | ΔBIC | Parameter estimates (95% CI) |
|---|---|---|---|
| $(\text{number of turns}) = \exp\,(a_0)$ | 611 | 27 | $a_0 = 0.0559\ (-0.0715\text{–}0.1834)$ |
| $(\text{number of turns}) = \exp\,(a_0 + D)$<br>$D = \begin{cases} 0 \text{ (in } 20-180 \text{ km from the goal)} \\ b \text{ (in } 180-500 \text{ km from the goal)} \end{cases}$ | 605 | 21 | $a_0 = 0.290\ (0.115\text{–}0.465)$<br>$b = -0.447\ (-0.702\text{–}-0.192)$ |
| $(\text{number of turns}) = \exp\,(a_0 + D)$<br>$D = \begin{cases} 0 \text{ (in } 20-340 \text{ km from the goal)} \\ b \text{ (in } 340-500 \text{ km from the goal)} \end{cases}$ | 596 | 12 | $a_0 = 0.205\ (0.0674\text{–}0.343)$<br>$b = -0.756\ (-1.21\text{–}-0.391)$ |
| $(\text{number of turns}) = \exp\,(a_0 + D)$<br>$D = \begin{cases} 0 \text{ (in } 20-180 \text{ km from the goal)} \\ b_1 \text{ (in } 180-340 \text{ km from the goal)} \\ b_2 \text{ (in } 340-500 \text{ km from the goal)} \end{cases}$ | 600 | 16 | $a_0 = 0.290\ (0.115\text{–}0.465)$<br>$b_1 = -0.210\ (-0.494\text{–}0.0732)$<br>$b_2 = -0.841\ (-1.22\text{–}-0.461)$ |
| $(\text{number of turns}) = \exp\,(a_0 + a_1\lvert\theta_P\rvert)$ | 612 | 28 | $a_0 = 0.353\ (0.151\text{–}0.555)$<br>$a_1 = -0.149\ (-0.307\text{–}0.00878)$ |
| $(\text{number of turns}) = \exp\,(a_0 + a_1\lvert\theta_P\rvert + D)$<br>$D = \begin{cases} 0 \text{ (in } 20-180 \text{ km from the goal)} \\ b \text{ (in } 180-500 \text{ km from the goal)} \end{cases}$ | 607 | 23 | $a_0 = 0.353\ (0.151\text{–}0.555)$<br>$a_1 = -0.149\ (-0.307\text{–}0.00878)$<br>$b = -0.430\ (-0.686\text{–}-0.174)$ |
| $(\text{number of turns}) = \exp\,(a_0 + a_1\lvert\theta_P\rvert + D)$<br>$D = \begin{cases} 0 \text{ (in } 20-340 \text{ km from the goal)} \\ b \text{ (in } 340-500 \text{ km from the goal)} \end{cases}$ | 599 | 15 | $a_0 = 0.353\ (0.151\text{–}0.555)$<br>$a_1 = -0.148\ (-0.304\text{–}0.00760)$<br>$b = -0.742\ (-1.11\text{–}-0.377)$ |
| $(\text{number of turns}) = \exp\,(a_0 + a_1\lvert\theta_P\rvert + D)$<br>$D = \begin{cases} 0 \text{ (in } 20-180 \text{ km from the goal)} \\ b_1 \text{ (in } 180-340 \text{ km from the goal)} \\ b_2 \text{ (in } 340-500 \text{ km from the goal)} \end{cases}$ | 602 | 18 | $a_0 = 0.428\ (0.202\text{–}0.654)$<br>$a_1 = -0.143\ (-0.299\text{–}0.0131)$<br>$b_1 = -0.197\ (-0.481\text{–}0.0867)$<br>$b_2 = -0.822\ (-1.20\text{–}-0.441)$ |
| $(\text{number of turns}) = \exp\,(a_0 + a_1\lvert\theta_P\rvert + a_2\lvert\theta_P\rvert^2)$ | 593 | 9 | $a_0 = 0.688\ (0.434\text{–}0.941)$<br>$a_1 = -1.48\ (-2.03\text{–}-0.937)$<br>$a_2 = 0.480\ (0.293\text{–}0.667)$ |
| $(\text{number of turns}) = \exp\,(a_0 + a_1\lvert\theta_P\rvert + a_2\lvert\theta_P\rvert^2 + D)$<br>$D = \begin{cases} 0 \text{ (in } 20-180 \text{ km from the goal)} \\ b \text{ (in } 180-500 \text{ km from the goal)} \end{cases}$ | 589 | 5 | $a_0 = 0.762\ (0.504\text{–}1.02)$<br>$a_1 = -1.35\ (-1.90\text{–}-0.795)$<br>$a_2 = 0.435\ (0.246\text{–}0.625)$<br>$b = -0.651\ (-1.02\text{–}-0.284)$ |
| $(\text{number of turns}) = \exp\,(a_0 + a_1\lvert\theta_P\rvert + a_2\lvert\theta_P\rvert^2 + D)$<br>$D = \begin{cases} 0 \text{ (in } 20-340 \text{ km from the goal)} \\ b \text{ (in } 340-500 \text{ km from the goal)} \end{cases}$ | 584 | 0 | $a_0 = 0.762\ (0.504\text{–}1.02)$<br>$a_1 = -1.35\ (-1.90\text{–}-0.795)$<br>$a_2 = 0.435\ (0.246\text{–}0.625)$<br>$b = -0.651\ (-1.02\text{–}-0.284)$ |
| $(\text{number of turns}) = \exp\,(a_0 + a_1\lvert\theta_P\rvert + a_2\lvert\theta_P\rvert^2 + D)$<br>$D = \begin{cases} 0 \text{ (in } 20-180 \text{ km from the goal)} \\ b_1 \text{ (in } 180-340 \text{ km from the goal)} \\ b_2 \text{ (in } 340-500 \text{ km from the goal)} \end{cases}$ | 587 | 3 | $a_0 = 0.852\ (0.574\text{–}1.13)$<br>$a_1 = -1.36\ (-1.91\text{–}-0.806)$<br>$a_2 = 0.441\ (0.252\text{–}0.630)$<br>$b_1 = -0.225\ (-0.509\text{–}0.0594)$<br>$b_2 = -0.742\ (-1.12\text{–}-0.360)$ |

Table S2 Results of fitting glm models to data on the number of turns made by the sailboats against the direction of the goal relative to the wind. The best model is shown in gray.

| Model | BIC | ΔBIC | Parameter estimates (95% CI) |
|---|---|---|---|
| $(\text{number of turns}) = \exp\,(a_0)$ | 151 | 36 | $a_0 = -0.965\ (-1.32\text{–}-0.614)$ |
| $(\text{number of turns}) = \exp\,(a_0 + D)$<br>$D = \begin{cases} 0 \text{ (in } 20 - 180 \text{ km from the goal)} \\ b \text{ (in } 180 - 500 \text{ km from the goal)} \end{cases}$ | 155 | 40 | $a_0 = -1.14\ (-1.80\text{–}-0.472)$<br>$b = 0.245\ (-0.537\text{–}1.03)$ |
| $(\text{number of turns}) = \exp\,(a_0 + D)$<br>$D = \begin{cases} 0 \text{ (in } 20 - 340 \text{ km from the goal)} \\ b \text{ (in } 340 - 500 \text{ km from the goal)} \end{cases}$ | 153 | 38 | $a_0 = -1.19\ (-1.67\text{–}-0.710)$<br>$b = 0.568\ (-0.134\text{–}1.27)$ |
| $(\text{number of turns}) = \exp\,(a_0 + D)$<br>$D = \begin{cases} 0 \text{ (in } 20 - 180 \text{ km from the goal)} \\ b_1 \text{ (in } 180 - 340 \text{ km from the goal)} \\ b_2 \text{ (in } 340 - 500 \text{ km from the goal)} \end{cases}$ | 157 | 42 | $a_0 = -1.14\ (-1.80\text{–}-0.472)$<br>$b_1 = -0.118\ (-1.08\text{–}0.849)$<br>$b_2 = -0.511\ (-0.328\text{–}1.35)$ |
| $(\text{number of turns}) = \exp\,(a_0 + a_1\lvert\theta_P\rvert)$ | 115 | 0 | $a_0 = 0.719\ (0.197\text{–}1.24)$<br>$a_1 = -3.60\ (-4.91\text{–}-2.30)$ |
| $(\text{number of turns}) = \exp\,(a_0 + a_1\lvert\theta_P\rvert + D)$<br>$D = \begin{cases} 0 \text{ (in } 20 - 180 \text{ km from the goal)} \\ b \text{ (in } 180 - 500 \text{ km from the goal)} \end{cases}$ | 120 | 5 | $a_0 = 0.675\ (-0.116\text{–}1.47)$<br>$a_1 = -3.59\ (-4.90\text{–}-2.28)$<br>$b = -0.0587\ (-0.726\text{–}0.844)$ |
| $(\text{number of turns}) = \exp\,(a_0 + a_1\lvert\theta_P\rvert + D)$<br>$D = \begin{cases} 0 \text{ (in } 20 - 340 \text{ km from the goal)} \\ b \text{ (in } 340 - 500 \text{ km from the goal)} \end{cases}$ | 119 | 4 | $a_0 = 0.574\ (-0.0725\text{–}1.22)$<br>$a_1 = -3.54\ (-4.85\text{–}-2.22)$<br>$b = 0.29\ (-0.420\text{–}1.00)$ |
| $(\text{number of turns}) = \exp\,(a_0 + a_1\lvert\theta_P\rvert + D)$<br>$D = \begin{cases} 0 \text{ (in } 20 - 180 \text{ km from the goal)} \\ b_1 \text{ (in } 180 - 340 \text{ km from the goal)} \\ b_2 \text{ (in } 340 - 500 \text{ km from the goal)} \end{cases}$ | 124 | 9 | $a_0 = 0.658\ (-0.139\text{–}1.45)$<br>$a_1 = -3{,}54\ (-4.86\text{–}-2.22)$<br>$b_1 = -0.167\ (-1.13\text{–}0.800)$<br>$b_2 = 0.208\ (-0.637\text{–}1.05)$ |

Table S3 Currency function estimates.

| Model | BIC | ΔBIC | Parameter estimates |
|---|---|---|---|
| Quadratic<br>$\mathcal{C}(\theta_G) = 1 + c_2(|\theta_G| - c_3)^2$ | -3,258 | 0 | $(\beta, c_2, c_3)$<br>$= (0.285, 0.314, 2.18)$ |
| Linear<br>$\mathcal{C}(\theta_G) = 1 + c_1|\theta_G|$ | -1,458 | 1800 | $(\beta, c_1) = (0.156, -0.164)$ |
| Constant cost<br>$\mathcal{C}(\theta_G) = 1$ | 343 | 3611 | $(\beta) = (0.187)$ |
